# Amino acid biostimulant increases radiata pine photosynthetic efficiency and growth through optimised mycobiome and nitrogen assimilation

**DOI:** 10.1101/2025.03.26.645424

**Authors:** Jamil Chowdhury, Nathan Milne, Melanie Wade, Robert Sharwood, Bronwyn Thuaux, Phil Green, Ian Last, John Senior, Angus J. Carnegie, Ian C Anderson, Stephen Elms, Krista L. Plett, Jonathan M Plett

**Affiliations:** Hawkesbury Institute for the Environment, Western Sydney University, Locked Bag 1797, Penrith, NSW, 2751, Australia; HVP Planations, Ti Tree Road, PO Box 10, Alberton Gunaikurnai, Vic 3971, Australia; Forestry Corporation NSW, Riverina Highlands Building 76 Capper Street, Tumut, 2720, Australia; HQPlantations, Nursery Rd, Toolara Forest QLD 4570, Australia; Forest Science, NSW Department of Primary Industries and Regional Development, Parramatta, NSW, 2150, Australia; Elizabeth MacArthur Agricultural Institute, NSW Department of Primary Industries and Regional Development, Menangle, NSW, 2568, Australia

## Abstract

**Background:** Amino-acid biostimulants have emerged as powerful alternatives to conventional inorganic nitrogen fertilisers, yet their potential in forestry species like radiata pine (*Pinus radiata*) remains largely unexplored. In this study, we reveal physiological mechanisms of enhanced growth of radiata pine seedlings that are achieved by substituting standard inorganic fertigation, either partially or entirely, with amino-acid-based biostimulants.

**Results:** Amino-acid fertigation notably increased shoot biomass, plant height, and collar diameter. Critically, this approach reshaped the root fungal community, selectively enriching fungi with diverse ecological roles, including several taxa known for auxin production. These microbial shifts correlated directly with elevated auxin concentrations observed in needle tissues, providing a plausible mechanism for the enhanced growth. Machine learning models further identified key fungal genera that strongly associated with plant biomass, reinforcing microbiome shifts as a contributing mechanism to enhanced growth. Additionally, amino-acid fertigation improved nitrogen assimilation, correlating positively with increased chlorophyll content and photosynthetic efficiency.

**Conclusions:** Our findings highlight that the transition from inorganic source to amino-acid biostimulants not only enhances plant growth and nitrogen use but also promotes a beneficial root microbiome, thereby offering a sustainable pathway to nursery production of radiata pine.

## Introduction

As one of the key macronutrients used by plants, nitrogen (N) is found in nearly all biomolecules that directly influence growth and productivity [1, 2]. Traditionally, in both agriculture and forestry, inorganic nitrogen fertilizers in the forms of nitrate (NO₃⁻) and ammonium (NH₄⁺), have been widely used due to their immediate availability and cost-effectiveness [3]. However, plants typically recover <50% of the applied fertiliser with the remainder being lost to leaching and denitrification [4]. Furthermore, excessive use of these fertilizers poses significant environmental risks including soil acidification and increased emissions of greenhouse gases, both of which can compromise soil health and biodiversity [5, 6]. As excessive inorganic fertiliser is putting the sustainability of our soils, and by extension our agricultural and forestry operations, at risk, new alternatives are needed.

In forestry, optimised N uptake in economically vital species like radiata pine (*Pinus radiata*) is crucial for enhancing biomass accumulation and ensuring sustainable timber production [7]. While global consumption of N fertiliser is 109 million tonnes/annum across all sectors, forestry typically minimises the use of fertilisers (e.g. 4kg ha^-1^ annum^-1^ vs 800 kg ha^-1^ annum^-1^ for horticulture [3]). This is due to economic considerations as well as the fact that pines are highly dependent on beneficial mycorrhizal fungi for sourcing growth-limiting nutrients and endophytic fungi that increase growth through modulation of plant hormone pathways, which can be negatively impacted at high N applications. The diversity of ectomycorrhizal fungi in pine forests has been identified as one of the leading factors driving productivity of these ecosystems [8, 9]. In these symbioses, the host root is colonised by an ectomycorrhizal fungus whose hyphae extend into the soil and source nutrients far beyond the root depletion zone [10–12]. These nutrients are then shared with the plant host in return for photosynthetically fixed carbon [13–16]. In endophytic fungal symbioses, several genera have been identified that can boost pine growth through the production of auxin (IAA) and gibberellic acid (GA)[17–19]. Both mycorrhizal and endophytic fungal associations can be disrupted and repressed should inorganic fertilisers be overused [20–23]. While this may be less of a concern in plantations due to historically low fertiliser use, to achieve a quality seedling prior to planting, growers use much higher fertilisation strategies in radiata pine nurseries. This is leading to an increasing interest in alternative N fertilisation strategies or sources that are both cost effective and more environmentally friendly [24].

Biostimulants have emerged as a promising alternative to replace or supplement inorganic N fertilisers[25, 26]. Biostimulants are categorized into seven main classes, with amino acids falling under the group of protein hydrolysates and other nitrogen-containing compounds. Collectively, these biostimulants enhance nutrient efficiency and crop quality, contributing to sustainable, low-input/high-output crop productions [27]. Amino acid fertilisers are designated as biostimulants of plant growth both directly through serving as a nitrogen source and as a signalling molecule influencing plant metabolic pathways, but also indirectly through alterations to the plant microbiome. In the vegetable and grains industries, amino acid-based fertilizers have been shown to promote growth through enhanced nutrient uptake efficiency and improved stress tolerance [28–30] Amino acid fertilisers have also been found to modulate physiological and biochemical processes including an increase in chlorophyll content that improve photosynthetic rates, and stimulate biomass accumulation in peanut, maize and lettuce [31][32, 33]. In addition, the interaction between amino acid fertilizers and root-associated microbial communities adds another layer of complexity. Addition of amino acid biostimulants is thought to drive microbial community assemblies to be more favourable to plant health. This can include promotion of root-associated fungi, including mycorrhizal associations and endophytes, that can mine organic N from soils and share it with a plant host to boost plant nutrient acquisition, hormone production, and overall plant health [34–36].

While the benefits of amino acid biostimulants have been documented in various crops, their applicability and efficacy to promote growth in forestry species like radiata pine are not well-established. Further, the pathways by which amino acid biostimulants may alter plant physiology to support plant growth, and how this correlates with root mycobiome diversity and function, are not well characterised in forestry species. Given that radiata pine evolved to grow in soils poor in inorganic N, and its dependence on the root mycobiome for establishment and growth in plantations settings [24, 37], we hypothesized that amino acid fertilisers have potential to be a more sustainable means of promoting plant growth by promoting plant physiological pathways (e.g. photosynthetic activity), as well as through promotion of their ectomycorrhizal communities.

To test our hypotheses and address this knowledge gap, we quantified a range of commercially relevant growth parameters in radiata pine seedlings treated with amino acid fertigation compared to conventional inorganic nitrogen fertigation. By integrating plant physiological measurements with root microbiome analyses and employing gradient boosting modelling to identify key microbial taxa influencing growth, this study provides a comprehensive understanding of how amino acid fertigation can enhance radiata pine seedling performance. Altogether, our findings have the potential to inform sustainable forestry practices, promoting the use of environmentally sustainable inputs that optimize plant growth while mitigating adverse ecological impacts.

## Methods

### Experimental design, materials and plant growth conditions

Radiata pine (*Pinus radiata*) seeds, collected from a Victorian seed orchard, were prepared according to the standard nursery strategy used by HVP Plantations (Victoria, Australia). Seeds were soaked in water for 24 hours, followed by cold stratification for 48 hours, then sown into nonsterile potting mix composed of composted pine bark and supplemented with slow-release fertilizer (composted pine bark; 2.5kg m^-3^ of 3-4 month slow-release fertiliser (NPK of 18:2.6:9.9), 5.5kg m^-3^ of 8-9 month controlled release fertiliser (NPK 17:2.3:10); Agsolutions Pty Ltd, Sydney, Australia). Seedlings were grown in Transplant Systems 45-cell forestry trays, with each cell containing approximately 93 cm^3^ of the potting mix. A starter fertigation (Peters Professional, 1 g/L) was applied to all seedlings for the first six weeks after germination. Beyond this period, seedlings received either a full nutrient inorganic fertigation (FN-IN, routine industry practice) or a full nutrient amino-acid fertigation (FN-AA). Both approaches supplied 400 mg/ seedling/week total nitrogen (in the form of NO₃⁻ and NH₄⁺) through an inorganic fertilizer (Seltec) or amino-acid fertilizer (Ful-fil Organic Amino Acid, Plantneeds.com), respectively. To ensure comparable nutrient profiles, the amino-acid treatment was supplemented with any additional elements found in the inorganic fertilizer. Half-strength variants (HN-IF and HN-AA) were also implemented by applying half of the total nitrogen amount. At five weeks after germination, a microbial inoculum derived from Australian pine forests and cultured on quarter-strength potato dextrose agar (PDA) was applied to all seedlings. The microbial species present in the inoculum are listed in Table S1. Fungicides AMISTAR® 250 SC (1000 ppm), Emblem (1000 ppm), and SWITCH® (160 ppm) were then rotated weekly from six to 20 weeks after germination. The experiment followed a randomized block design with six replicate blocks for each treatment, and each block contained five seedlings, for a total of 30 seedlings per treatment. All plants were grown for six months in a controlled-environment chamber at the Hawkesbury Institute for the Environment, under a 12 h light/12 h dark cycle at 25 °C and 70% relative humidity, with illumination at approximately 700 LUX.

At harvest, plant height (PH) was measured by measuring the stem length from the soil surface to the top of the crown. Root collar diameter (CD) was the stem diameter at the soil surface. Potting mix was gently removed from the root ball using clean running water, and root colonization (MYC_avg) was measured by counting an average of 100 lateral roots per plant from randomly selected areas of the root system using a Zeiss stereomicroscope and classifying them as either colonised (i.e. the presence of short roots showing a formation of a mantle-like structure) or uncolonized and the percentage colonisation by ectomycorrhizal fungi calculated. Shoot and roots were separated and oven dried at 42°C for 4 weeks before measuring the dry shoot and root biomass (SW and RW, respectively). Shoot N content was analysed on ground shoots at Brookside Laboratories (New Bremen, Ohio, USA).

### Measurement chlorophyll fluorescence parameters and needle chlorophyl content

Chlorophyll fluorescence parameters were measured using a handheld multi-spectrometer (MultispeQ V 2.0, PhotosynQ.com). To ensure full coverage of the light guide, 5–6 needles were arranged horizontally under a custom-made light guide mask, as recommended by the manufacturer (https://help.photosynq.com/instruments/light-guide-mask.html#make-the-mask). While chlorophyll fluorescence-based measurements do not require the leaf to fully cover the light guide, as we found a fully covered light guide appears to provide more consistent measurements than partially covered light guide, we proceeded with this approach. Photosynthesis-related parameters were measured according to Photosynthesis RIDES 2.0 protocols (https://photosynq.org/protocols/photosynthesis-rides-2-0) with default instrument settings. Various chlorophyll fluorescence parameters were then compared to the full-nutrient inorganic fertilizer (FN-IF) control by calculating effect sizes (Glass’s Δ), and these results were presented as radar plots created by R (version 4.2.3). Immediately following fluorescence measurements, needles were snap-frozen in liquid nitrogen and stored at −80 °C for subsequent chlorophyll extraction. Representative samples of three to five needles (first true leaves) were ground in 1 mL of ice-cold methanol (100%) with a bead basher, following the methanol extraction method described by [38].

The resulting extracts were analyzed in a microplate reader, and absorbance values were corrected to a 1-cm pathlength. Chlorophyll concentrations were then calculated according to [38, 39] using the equations:

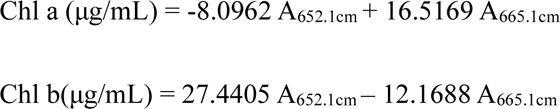

Finally, chlorophyll content was normalized to tissue weight.

### Hormone extraction and profiling

For the hormone profiling, first true needles (6-7) were harvested from each sample, immediately snapped into liquid nitrogen and stored at −80°C until hormone extraction. We used the Aqueous Phase 1 fraction for the hormone profiling which was extracted by a modified 3 pH Bligh-Dyer extraction procedure [40]. In brief, around 100 mg (equivalent to 0.8 parts of the extraction procedure) of needles were ground by an MPBio bead mill (6 Hz, 2x 30sec each) followed by adding 2 parts of the ice-cold methanol and sonicating to make sure that the tissue is properly homogenised. Following sonication, 1 part of chloroform was added, incubated in ice for half an hour. After that we added 1 part of water to each sample followed by adding 1 part of chloroform. Samples were vortexed each time immediately after adding each liquid. In the final step, we centrifuged the samples for 30 minutes at 4°C at 5,000 ppm. We carefully saved and dried down the upper aqueous phase 1 and stored in −80°C before reconstituted in water for hormone profiling. Hormone profiling was performed with an UltraPerformance liquid chromatograph (Sciex 7500 QTrap) by the Mass Spectrometry Facility of Western Sydney University.

### Statistical analysis of morphological and physiological data

To assess the statistical significance of each treatment effect on growth, photosynthetic and root colonization parameters, we calculated two-tailed p-values based on a t-statistic derived from the difference between control (FN-IF) and treatment groups, assuming pooled variance and (n1 + n2 − 2) degrees of freedom. These p-values were then compared against α = 0.05, and corresponding confidence intervals (75%, 85%, and 95%) were computed using this t distribution framework. To quantify the practical significance of the treatment, we calculated Glass’s Δ as the effect size measure which normalizes the difference between the means of the treated and control groups to the standard deviation of the control group. The choice of effect size measure, calculation and effect categorization were performed based on Maher et al. (2013) [41]. Pairwise correlation coefficients were calculated via Pearson’s r for each pair of variables, and p-values were derived from t-statistics to assess statistical significance. The resulting lower-triangle correlation matrix was visualized as a heatmap, with p-value thresholds used to annotate or highlight significant relationships. A correlation network was then constructed by filtering edges for correlations with p < 0.05 and mapping them in an undirected graph, where node position and edge width/color reflect the strength and direction of the significant correlations. All the analyses were performed using R (version 4.2.3).

### DNA extraction, MiSEQ library preparation, and Next generation sequencing

For DNA extraction, lateral roots taken randomly throughout the root system of three plants per block replicate were collected, pooled, and frozen. In total, six separate block replicates per treatment were extracted and prepared for community sequencing (n=6). DNA was extracted from approximately 100 mg of root sample using the ISOLATE II Plant DNA Kit (Meridian Bioscience) following manufacturer’s protocol. A two-step PCR protocol for Illumina MiSeq Meta-barcoding was followed to amplify the ITS region of fungi as described by [42] using 20 ng of the extracted gDNA and the following primers to amplify the ITS2 region (Forward: 5’-CCTACACGACGCTCTTCCGATCTNNNNGTGARTCATCGAATCTTTG-3’ and Reverse: 5’-GTGTGCTCTTCCGATCTCCTCCGCTTATTGATATGC-3’). Samples were multiplexed and sequenced on the Illumina MiSeq platform at the Advanced Gene Technology Centre, New South Wales Department of Primary Industries and Regional Development. DNA extraction blanks controls were processed in parallel with samples. All controls produced < 200 raw reads and < 10 post-filter reads, and shared no unique ASVs; consequently they were excluded from the final phyloseq object.

### Raw sequence processing and microbial population analysis

Demultiplexed paired-end raw sequences, as obtained from the sequencing facility, were processed by DADA2 [43]. The amplicon workflow was implemented by the DADA2 R package which include filtering, dereplication, sample inference, chimera identification, and merging of paired-end reads. The DADA2 R package was also used to assign the fungal taxonomy to the inferred ASVs with UNITE database (dynamic release v9, release date 2023-07-18, DOI: 10.15156/BIO/2938066) [44]. The processed data was merged with sample metadata creating a single object with Phyloseq R package [45] for analysing and visualization with relevant R packages in R version 4.2.3). Datasets were rarefied to 10 000 reads per library using “phyloseq::rarefy_even_depth()” before calculating Shannon indices with “estimate_richness()”. Beta diversity was quantified via Bray–Curtis distances and visualized using Principal Coordinates Analysis (PCoA) [46] [47]. Read counts at the genus level were normalized to per-sample totals, resulting in fractional abundances that sum to 100% per sample. These normalized values were then aggregated by treatment group to facilitate comparisons. Differential abundance analyses were conducted at the genus level with DESeq2 [48]. Phyloseq objects were aggregated to genus, and raw counts were transformed into a DESeq2 object specifying the main experimental factor (Fertilizer treatment) in the design formula. DESeq2 was then run to detect genera with significant log₂ fold changes in abundance (adjusted p < 0.05). Genus names were retrieved from UNITE taxonomic database, and significantly changing taxa were visualized in bar plots ordered by their log₂ fold change. Genus-level co-occurrence networks were inferred with SPIEC-EASI (mb method, λ.min.ratio = 1×10⁻3, nlambda = 60, pulsar rep.num = 50, StARS thresh = 0.05) on CLR-transformed relative-abundance matrices [49, 50]. Adjacency matrices output by SPIEC-EASI were converted into undirected graphs using the igraph R package(igraph package;[51]), and network topology metrics (e.g., node degree, clustering coefficient, and modularity) were computed. Community detection was performed via fast-greedy or similar algorithms to identify modules and hub taxa. Fungal lifestyles were annotated by matching genus-level taxa to entries in a FungalTraits database [52], which classifies fungal genera into categories such as ectomycorrhiza, saprotroph, pathogen, symbiont, unknown, etc. This lifestyle annotations were integrated with relative abundance data and differential abundance data.

### Machine learning approaches on microbial population analysis

To predict the impact of key microbial genera on plant growth performance, we employed a Gradient Boosting Machine (GBM) model [53], with shoot dry biomass as the metric. Plants with shoot dry biomass above 1.75 grams were categorized as “Healthy Growth” whereas those with lower biomass were classified as “Poor Growth.” The growth condition data was then combined with this matrix for further analysis. To train and validate the model, we divided the dataset into training and testing sets using the “createDataPartition” function from the “caret” package [54]. The training set consisted of 80% of the data, while the remaining 20% was allocated for testing. The training set was used to build the model, with the testing set reserved for performance evaluation. The model was subjected to 10-fold cross-validation, repeated three times, to minimize overfitting and ensure robustness. We evaluated the model primarily by the area under the ROC curve (AUC). To optimize the GBM’s hyperparameters (number of trees, interaction depth, learning rate, and minimum observations in terminal nodes) a grid search was conducted using the “train” function from the “caret” package. We proceeded to determine the relevance of each genus in predicting growth conditions by utilizing the “varImp” function. This analysis allowed us to identify the top 20 genera based on their overall importance scores. We examined the effect of these top genera on growth conditions using Partial Dependence Plots (PDPs) created with the “iml” package [55]. These plots reveal the marginal effect of individual genera on the likelihood of “Healthy Growth.” We evaluated the GBM model’s performance by generating confusion matrices and ROC curves for both training and testing datasets, using the AUC to measure the model’s ability to discriminate between healthy and poor growth. Additionally, we refined our analysis by quantifying the impact of each genus on healthy plant growth. This involved calculating the difference between predicted probabilities at the start and end points of each genus’s abundance. Finally, the results were visualized in a horizontal bar plot, clearly showing whether each genus contributed positively or negatively to healthy growth.

## Results

### Applying N as amino acids significantly increase shoot biomass of radiata pine compared to equivalent levels of inorganic N

The application of N in the form of amino acid (AA) fertilizers had significant effects on the growth and development of radiata pine seedlings (Fig1, Fig S1, Table S2). We observed significant changes in several key morphological traits across treatments. Shoot biomass, when comparing industry routine FN-IF to both FN-AA and HN-AA, increased >40% and 35%, respectively (Fig. 1B). The effect sizes were statistically significant, and when evaluated against the benchmarks established by [56] and [57], they were classified as ‘Large’ and ‘Very large’, respectively. The effect size of FN-AA fertigation on plant height was ‘small’, but significant (p=0.013), showing an average 10% increase over FN-IF. Root biomass decreased significantly in the HN-IF (11%), FN-AA (20%), and HN-AA (9%) treatments, although the effect size was only significant in the FN-IF vs. HN-AA (p=0.002). Altered allocation of growth was especially strong in root:shoot ratio where AA-treated plants showed dramatic reductions in this ratio of -46% for FN-AA and -35% for HN-AA, when compared to FN-IF. The effect size was highly significant (p<1.73×10^-10^), indicating a very large impact of the fertigation regime on this parameter. No significant difference in the percentage of roots exhibiting mycorrhizal colonisation was observed between the treatments.

**Figure 1:**
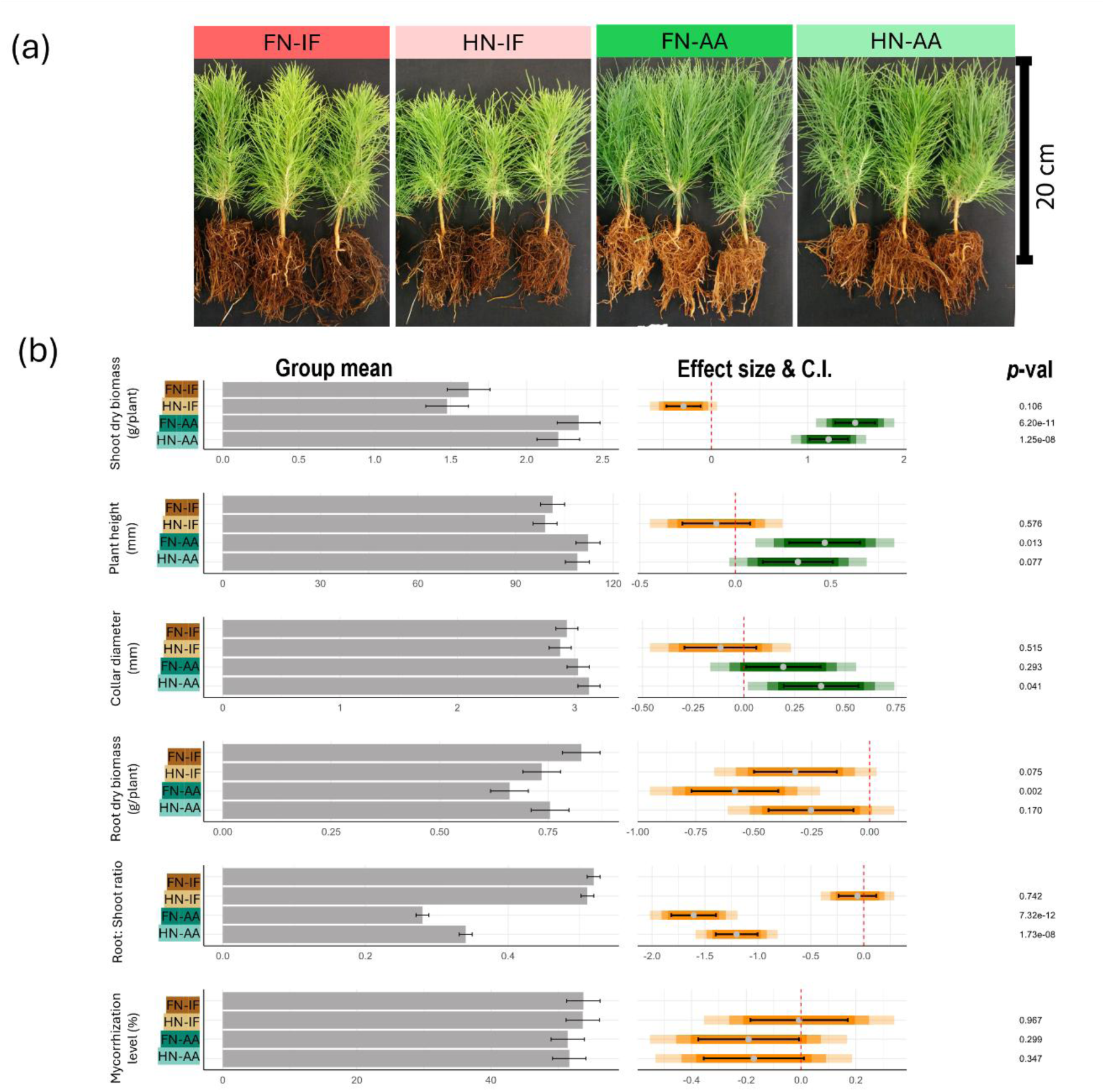
Impact of amino acid fertigation on the morphology of radiata pine seedlings. (a) Photographs of seedlings treated with various form and level of fertigation. (b) left panel presents group means for each variable, with error bars indicating standard errors. In the right panel, coloured bars represent effect sizes (Glass’s Δ), with grey dots marking the mean effect of each fertilizer type relative to the control fertigation (FN-IF). The coloured bars depict confidence intervals (C.I.) at 95%, 85%, and 75% levels, where lighter shades indicate higher confidence intervals. If a coloured bar intersects the zero reference line (red dotted line), the fertilizer effect is not statistically different from the control at the respective confidence level. Black bars denote the standard error of the mean (SEM) for each treatment. Green and orange bars indicate positive and negative effects of the treatments, respectively. FN-IF = Full nutrient ingorganic fertigation, HN-IF = Half nutrient inorganic fertigation, FN-AA = Full nutrient amino-acid fertigation, HN-AA= Half nutrient amino-acid fertigation.

### Amino acid fertigation enhances photosynthetic efficiency and modulates energy dissipation in radiata pine

The use of AA fertilizers had a significant impact on needle chlorophyll content and total nitrogen levels (Fig 2, Table S3). Both FN-AA and HN-AA increased chlorophyll-a content by 50%, while FN-AA increased chlorophyll-b by around 40% and HN-AA by nearly 30%. Additionally, the application of FN-AA increased total nitrogen levels in the needle by around 50%, and HN-AA increased it by around 40%. These findings are further supported by observations of the chlorophyll fluorescence parameter, SPAD, which increased by more than 10% in amino acid-treated plants (Fig. 2B).We also observed a minimal negative effect on total nitrogen content and chlorophyll levels when the control fertigation (FN-IF) was reduced to half (HN-IF). Due to the increased level of nitrogen and total chlorophyl content in amino-acid treated seedlings, we measured photosynthetic efficicency of the needles. Plants treated with amino acid-based fertigations ((HN-AA and FN-AA) exhibited small but significantly positive effect on the efficiency of Photosystem II (PSII), as evidenced by increased Maximum Quantum Efficiency (Fv/Fm) and Quantum Yield (ΦPSII) [58]. The Fv/Fm ratio showed a 4.1% increase under FN-AA treatment and a 3.8% increase under HN-AA. Amino acid fertigation also positively influenced Photosystem I (PSI) parameters with a very large positive effect in PS I active centres (PSI_ACT) and the PS I over-reduced centres (PSI_OR) states (increased by 53% and 157% respectively; Fig 2B, Table S4). The improvements in both PSII and PSI efficiency under amino acid treatments were further supported by the observed increase in Linear Electron Flow (LEF). Plants treated with inorganic fertigation (HN-IF and FN-IF), in contrast, demonstrated a greater reliance on protective energy dissipation mechanisms, as indicated by increased non-photochemical quenching (NPQ)-related parameters (ΦNPQ and NPQt; Fig 2B)

**Figure 2:**
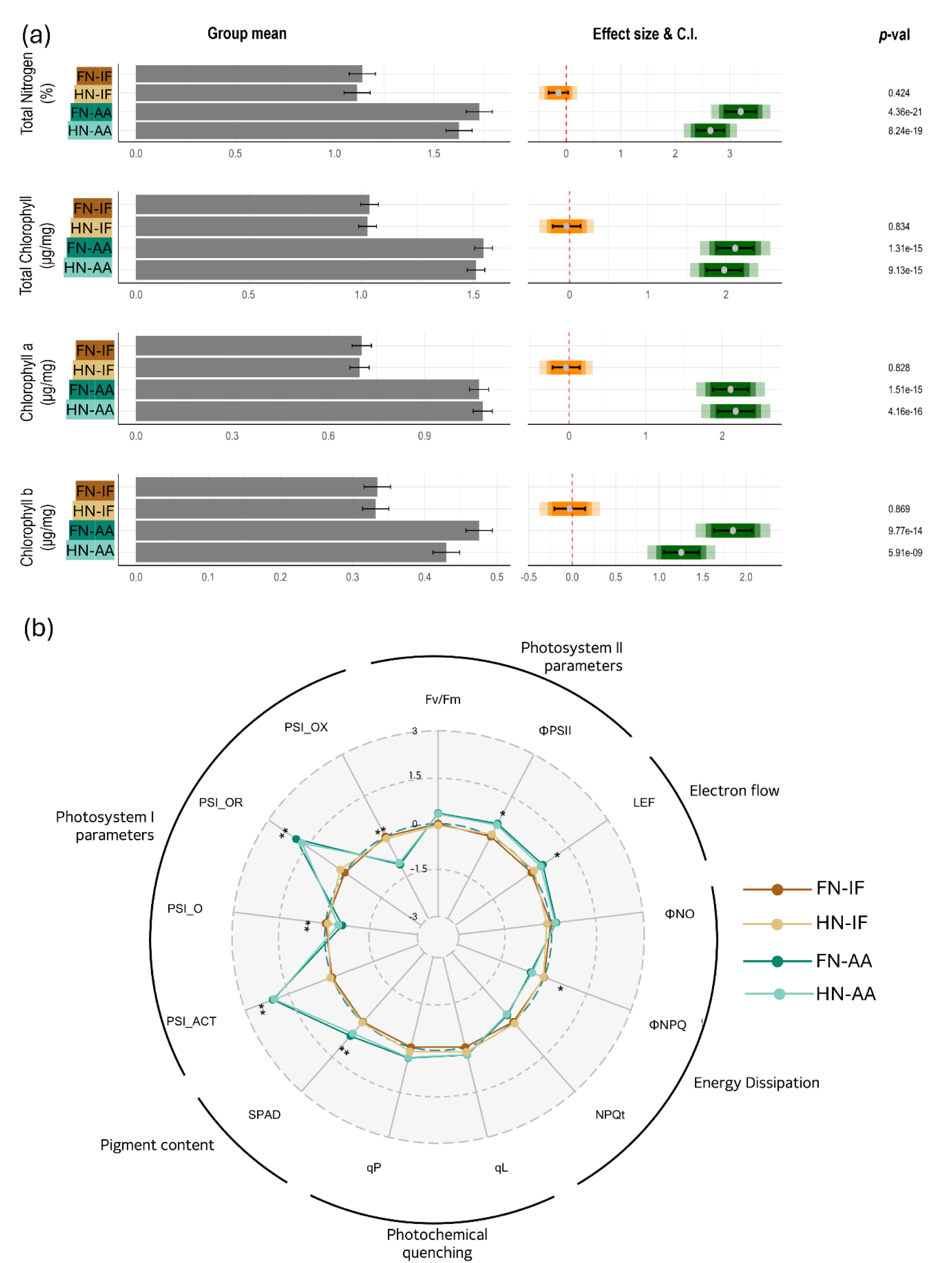
Effect of amino acid fertigation on needle chemistry and photosynthetic activity. (a) In the left panel, bars shows the group mean of the respective variable with standard error. In the right panel, color bars shows effect size (Glass’s Δ) where the grey dots indicates the mean effect of respective fertiliser types compared to control fertigation FN-IF. The colored bars indicate the confidence intervals at 95%, 85%, and 75% levels (ligher color inidcate higher C.I.). If the color bar intercepts with zero (red dottted line) it means that the fertizer effect is not statistically significant from control at the respective confidence interval. Black bars indicate SEM values of the treatment mean. (b) A radar plot shows the effects of fertilizers on selected chlorophyl fluorescence parameters. The value of each parameter represents the effect size (Glass’s Δ) calculated against control treatement (FN-IF). ** indicates p-value <0.01, * indicates p-value < 0.05.

### Root fungal diversity and network significantly altered by changing fertigation from inorganic to organic amino acid form

Amino acid-treated plants had significantly increased fungal diversity in radiata pine root system as measure by Shannon diversity index [59] (Fig. 3A), although there was no significant difference between FN-AA and HN-AA. Despite the observed increase in richness, the evenness of the fungal communities as measured by Pielou’s Evenness Index [60] remained largely unaffected by the amino acid fertigation (Fig. 3B). Bray-Curtis Principal Coordinates Analysis (PCoA) revealed distinct clustering patterns based on the type of fertilizer applied (Fig. 3C). The top 20 fungal genera displayed a significant variation in their relative abundance depending on the fertigation type (Fig. 4A). A substantial proportion of these genera possess an unknown lifestyle (Fig. 4B). While well-known pine ectomycorrhizal fungi from North American and European samples were notably absent from the top 20 genera list, several species belong to ectomycorrhizal groups of genera *Sistotrema* [61] and *Serendipita* [62] were present. The relative abundance of these genera was notably reduced in seedlings treated with amino acid fertigation as opposed to inorganic fertigation. The genus *Pseudoarthrographis*, which has been characterized as plant pathogen in FungalTraits database [52], dominated the fungal community in plants treated with inorganic fertigation. In contrast, *Xenoacremonium*, a wood saprotroph, was more prevalent in plants treated with amino acid fertigation. Another significant observation was the consistent presence of *Meliniomyces* across all fertigation types. While this genus is typically characterized as root-endophyte [52], it can also form ectomycorrhiza [63]. Additionally, *Fusarium*, a well-known genus containing pine root rot pathogens [64], was commonly found across all fertigation treatments.

**Figure 3:**
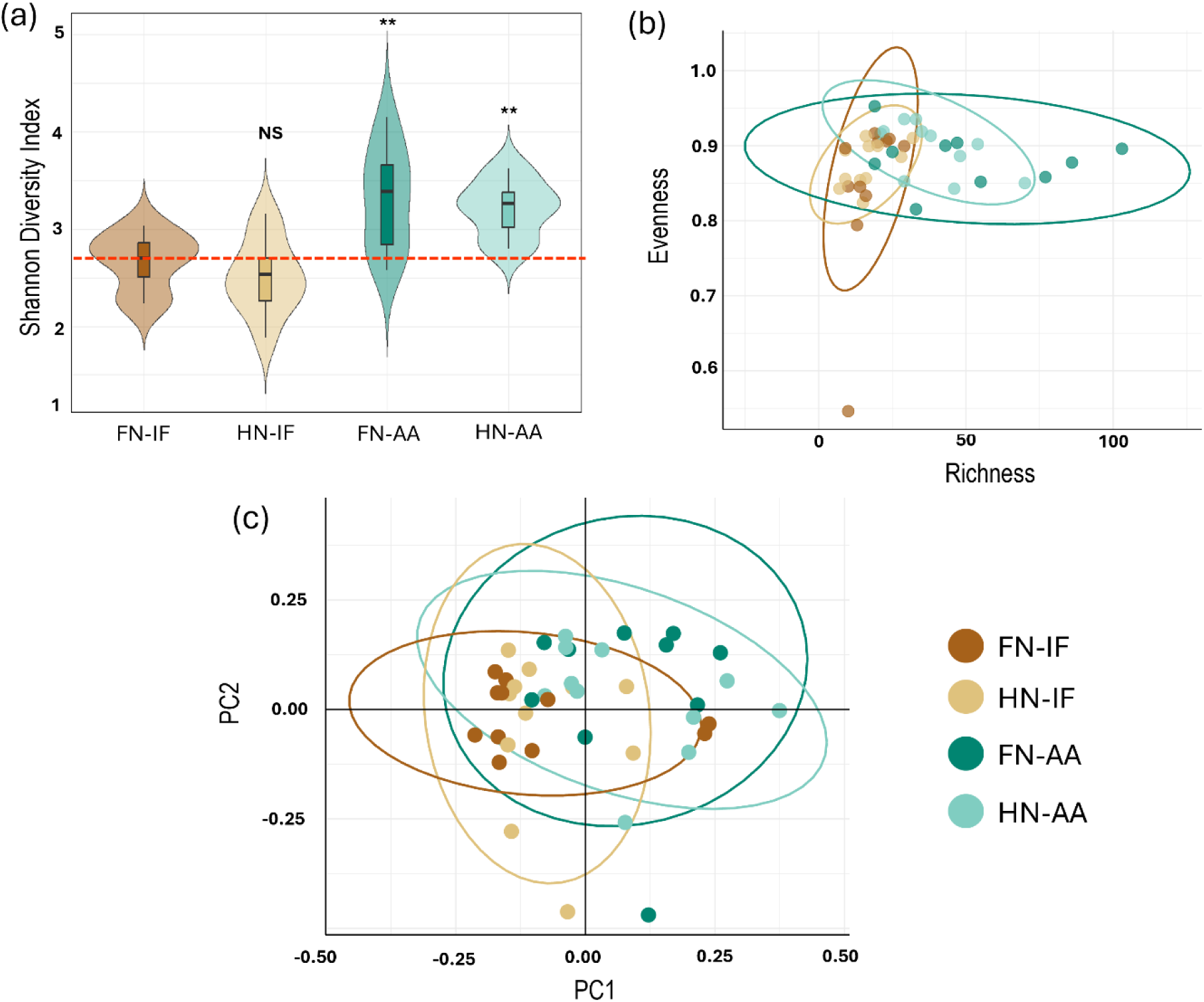
Impact of fertigation on root fungal diversity in radiata pine seedlings. (A) Shannon Diversity Index across different fertilizer treatments. The double asterisks (**) indicate significant differences (p < 0.01) between the treatments, as determined by ANOVA followed by Tukey’s post-hoc test. NS indicates non-significant differences. Red dotted line indicates the mean diversity index of control treatment (FN-IF) (B) Scatter plot of species richness (x-axis) and the evenness index (y-axis) is plotted. (C) Principal Coordinates Analysis (PCoA) of Bray-Curtis distances, showing beta diversity among the different fertigation treatments. The first three principal coordinates are plotted. Each point represents a sample, and the clustering patterns illustrate the compositional differences in fungal communities based on fertigation type. Colors correspond to the same treatment groups for all the plots. FN-IF= Full nutrient inorganic fertigation, HN-IF= Half nutrient inorganic fertigation, FN-AA= Full nutrient amino acid fertigation, HN-AA = Half nutrient amino acid fertigation.

**Figure 4:**
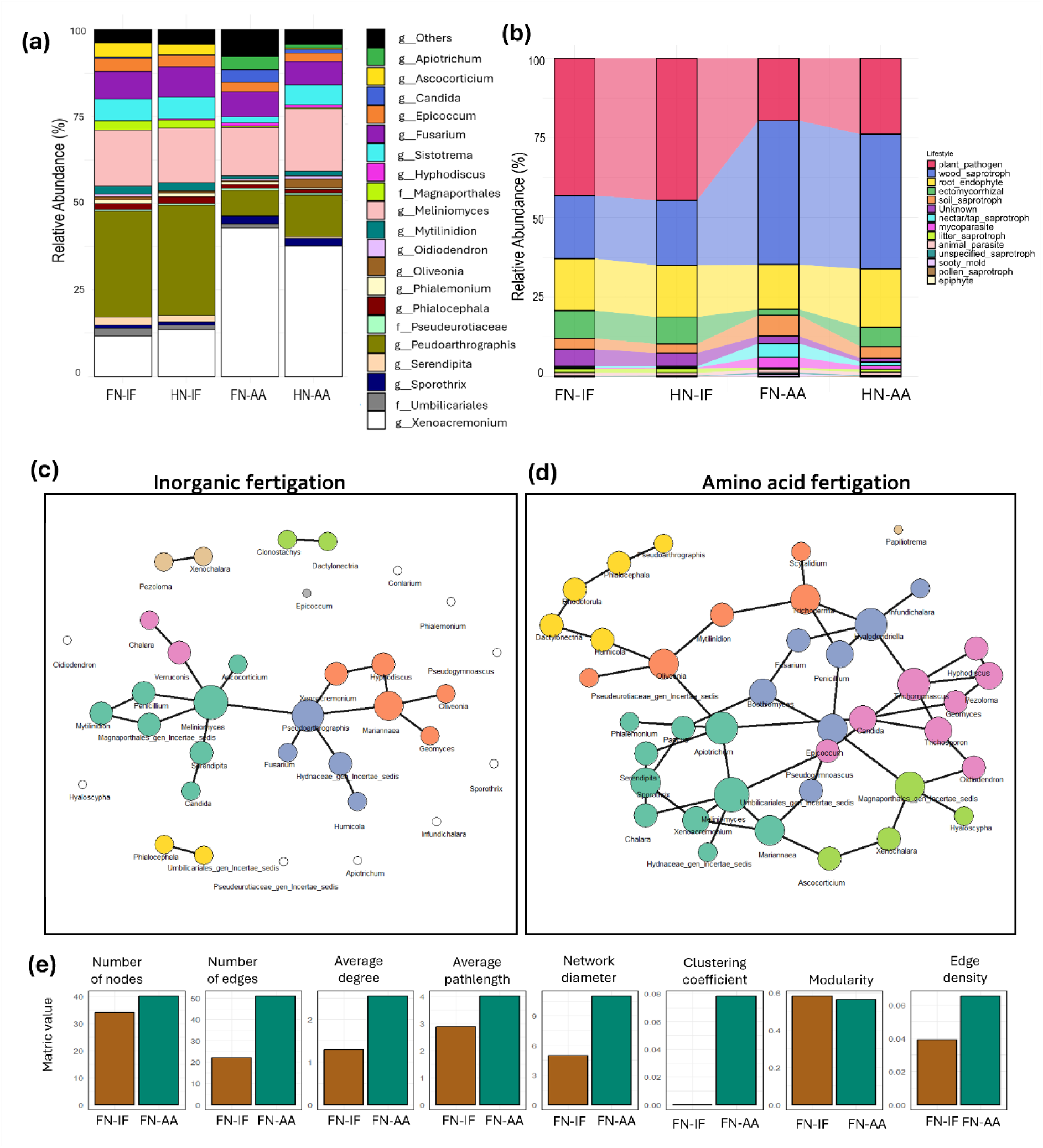
Relative abundance and fungal community structure as affected by fertigation. (a) Relative abundance of the top 20 genera. (b) Functional lifestyle profile (relative abundance) predicted by FungalTraits [52]. Co-occurrence networks of fungal communities in Radiata pine roots under inorganic (C) and amino acid fertigation (D). Each node represents a fungal genus, with node size proportional to degree centrality. The same color in nodes represents genera within the same module, indicating closely interacting groups.

Co-occurrence network analysis (Fig. 4C,D,E) revealed clear structural and compositional differences in the fungal communities associated with radiata pine roots under amino acid vs. inorganic fertigation regimes (Figure 4B,C). Under amino acid fertigation (AA), the co-occurrence network consisted of 40 nodes with 51 edges. The AA network also exhibited a diameter of 11, an average path length of approximately 4.01, and a global clustering coefficient of around 0.08, indicating that many taxa formed tightly interconnected clusters yet were still reachable from one another within just a few steps. The AA network showed a lower edge density (∼0.065), implying that despite having more nodes and edges, it was proportionally more “spread out” among its potential connections. By contrast, the inorganic fertigation (FN-IF) network was more compact, with 34 nodes with 22 edges. It had a smaller diameter (5) and lower edge density (∼0.039), reflecting a tighter proportion of connections among fewer taxa.

The clustering coefficient (0) and average path length (∼2.89) suggested that community members in this network also formed closely linked clusters, though with fewer overall participants. Notably, both networks exhibited substantial compartmentalization (as evidenced by their moderately high modularity values), suggesting that each fertigation regime selects for distinct fungal “modules” or subcommunities. When comparing the key hub genera, the amino acid fertigation network featured *Apiotrichum*, *Oliveonia*, *Trichoderma*, *Hyalodendriella*, and *Trichomonascus* as the top hubs. In contrast, the hub genera in the inorganic fertigation network were *Meliniomyces*, *Pseudoarthrographis*, and *Mariannaea*, each demonstrating high connectivity and importance in structuring the fungal community under this treatment. These results highlight the significant impact of fertigation type on fungal community structure, with amino acid fertigation supporting a larger and more compartmentalized fungal community, whereas inorganic fertigation yielded a smaller but relatively denser community of connected taxa.

### Differential recruitment of auxin-producing and pathogenic fungi in response to amino-acid fertigation in radiata pine

To identify specific genera that were significantly enriched or depleted under amino-acid fertigation, including minor genera, we performed differential abundance analysis comparing microbial communities in amino-acid-treated and inorganic-fertilized plants. This analysis identified 34 taxa that showed significant differences in abundance between treatments (Fig. 5). Amino-acid fertigation specifically enriched several fungal genera known to produce the phytohormone indole-3-acetic acid (IAA). Key auxin-producing fungi significantly enriched under amino-acid fertigation included *Trichoderma* [65, 66], *Apiotrichum* [67], *Rhodotorula* [68], *Papiliotrema* [69], *Candida* [70], and *Lecanicillium* (previously known as *Verticillium*) [71]. Additionally, isolates belong to *Rhodotorula* were also found phosphate solubilizing [72] and isolates belong to *Papiliotrema* are shown to improve nitrogen nutrition in plants [73]. The amino-acid treatment led to a notable reduction in the abundance of key ectomycorrhizal fungi, including *Serendipita* and *Sistotrema*. We also found a decrease in the abundance of certain pathogenic fungi, such as *Dactylonectria*, a genus known for causing root-associated diseases in conifers [74, 75].

**Figure 5:**
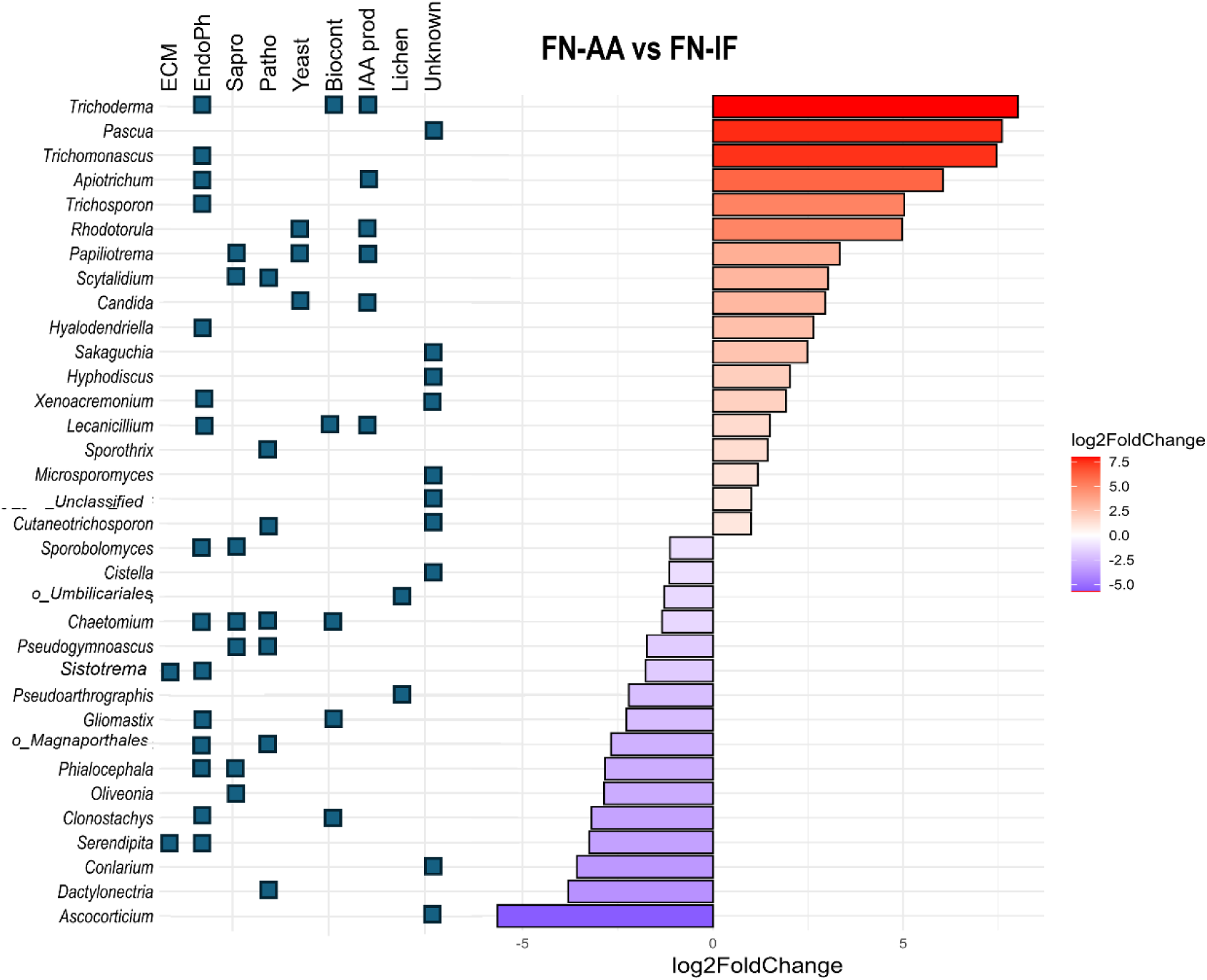
Differentially abundant genera in pine seedling roots treated with amino-acid fertigation compared to inorganic fertigation. The comparison was done only between FN-AA and FN-IF. Log2 fold change values are shown. Only the genera above the threshold (p-value <0.05, Log2FC=1) are shown. Known lifestyles belonging to these genera are indicated with blue boxes. ECM= Ectomycorrhizal, Endoph = Endophyte, Sapro = Saprotroph, Patho = Pathogen, Biocont. = Biocontrol agent, IAA prod = Fungi associated with IAA production, Lichen = Fungi associated with lichen formation.

### Amino-acid fertigation alters needle hormone profiles in a manner correlating to shoot biomass gains

Given the number of fungal genera that produce phytohormones showing differential abundance in amino acid treated plants versus inorganic N, we assayed the phytohormone profile of radiata pine needles. Notably, the levels of key growth-regulating hormones including indole-3-acetic acid (IAA), abscisic acid (ABA), gibberellic acids (GAs), and salicylic acid (SA) were influenced by the fertigation treatments (Fig 6, Table S5). In this particular scenario, we compared FN-AA with FN-IF treatments, and found that IAA levels exhibited a modest yet statistically significant increase of 13% under amino-acid fertigation (p=0.021) while ABA increased by 40%, a nearly significant result (p=0.053). This observation is consistent with the previous studies that amino acid-biostimulants, which contain a mixture of various amino-acids, can elicit auxin-like activities in shoot [76, 77]. Meanwhile, GA_3_, GA_4_, and JA-Ile, the biologically active forms of these hormones, were significantly repressed (Fig 6). Correlation network analysis showed a positive relationship between IAA levels and shoot biomass/collar diameter, while JA was in opposition to these traits (Fig 6B). Shoot nitrogen was negatively correlated to GA_3_, but was positively correlated to microbial diversity. A correlation matrix between major fungal genera present in pine seedlings and needle phytohormones is presented in tableS6.

**Figure 6:**
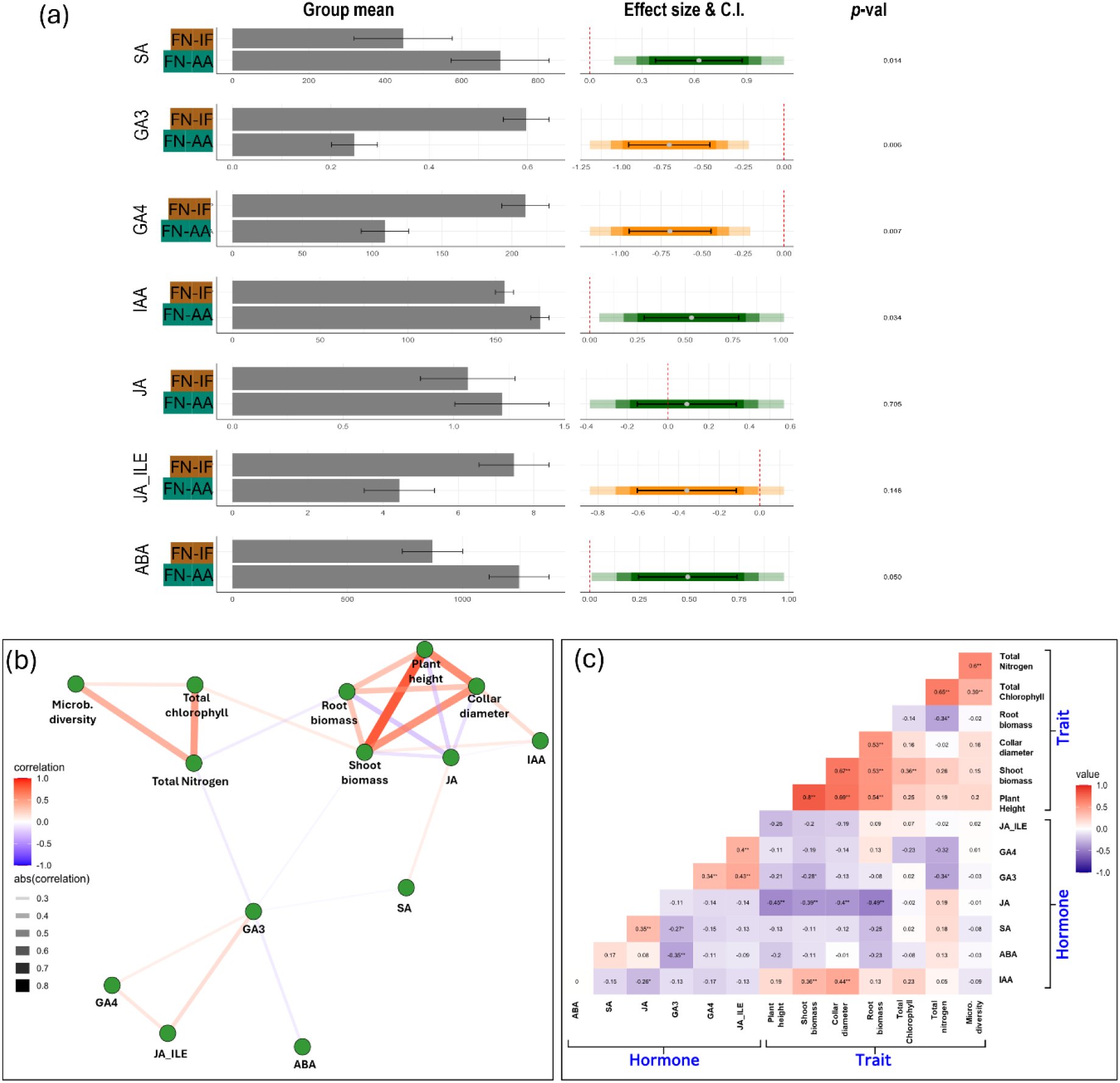
Effect of aminoacid fertigation on needle phytohormone level of radiata pine seedling. In the left panel, bars shows the group mean of the respective variable with standard error. In the right pannel, color bars shows effect size (Glass’s Δ) where the black dots indicate the mean effect of respective fertiliser types compared to control fertigation FN-IF. The colored bars indicate the confidence intervals at 95%, 85%, and 75% levels (ligher color inidcate higher C.I.). It the color bar intercepts with zero (red dot line) which means that the fertizer effect is not statistically significant from control at the respective confidence interval. Black bars indicate SEM values of the treatment mean. Pearson correlation between phytohormone and morphological traits of radiata pine seedlings. (A) Correlation network revealing the significant relationship among the variables. Line thickness indicates strength of the relationship between the connected parameters. Color and intensity represents Pearson’s R value which indicates of color direction and strength of the relationship. A threshold of p-value <0.05 is used. (B). Correlation matrix plot showing the strength of relationship with p-value among all the variables.

### Predictive modelling of fungal community contributions to radiata pine plant performance using Gradient Boosting

Given the positive correlation of microbial diversity to chlorophyll content and, thus, to plant biomass, we employed a Gradient Boosting Machine (GBM) model to quantify the contribution of specific fungal taxa to plant biomass. The model’s predictive performance was robust, achieving an accuracy of 93% and an area under the ROC curve (AUC) of 0.983 on the training dataset (Fig 7A). On the test set, the model retained good accuracy (71%) and a reasonable AUC (0.802), indicating high reliability in predicting plant growth outcomes based on root microbiome composition (Fig 7B). The model identified *Xenochalara*, *Apiotrichum*, and *Meliniomyces* as the top three genera contributing most significantly to shoot biomass (Fig 7C).

**Figure 7:**
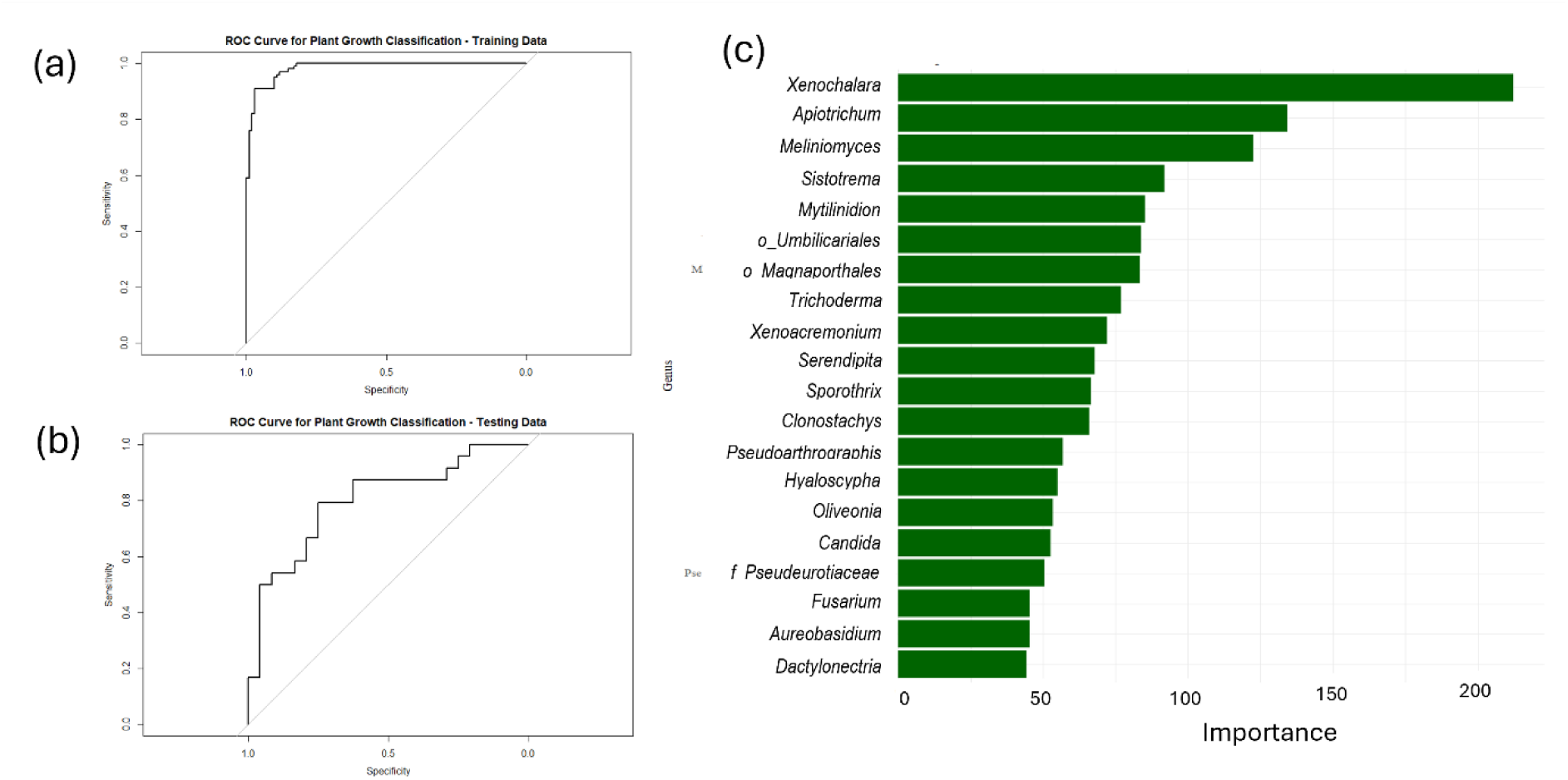
Gradient boosting model results for predicting plant biomass based on microbiome composition. (a) Top 20 most influencial fungal genera contributing to plant biomass. Importance is measured by the contribution of each genus to the prediction of Healthy or Poor shoot biomass at a given threshhold (1.75g dry weight). (b) ROC curve for the classification of plant growth conditions based on the training dataset, showing high predictive performance (AUC = 0.983). (c) ROC curve for the classification of plant growth conditions on the testing dataset, demonstrating a reasonable predictive accuracy (AUC = 0.802).

Partial dependence plot (PDP) analysis revealed that taxonic features such as *Apiotrichum*, *Aureobasidium*, *Candida*, *Sistotrema*, *Xenoacremonium*, and *Meliniomyces* positively contributed to healthy biomass (Fig. 8). Conversely, several taxa were associated with a reduction in shoot biomass, particularly those with saprotrophic or pathogenic lifestyles. Among these, *Dactylonectria, Xenochalara, Sporothrix, Oliveonia,* and several unclassified taxa such as *Pseudeurotiaceae, Magnaporthales,* and *Umbilicariales* were identified as contributing to poor plant growth (Fig. 8). A third pattern emerged for genera like *Fusarium*, *Serendipita*, and *Mytilinidion*, showing positive contributions to biomass at lower abundance levels but negative effects when their abundance exceeded a certain threshold.

**Figure 8:**
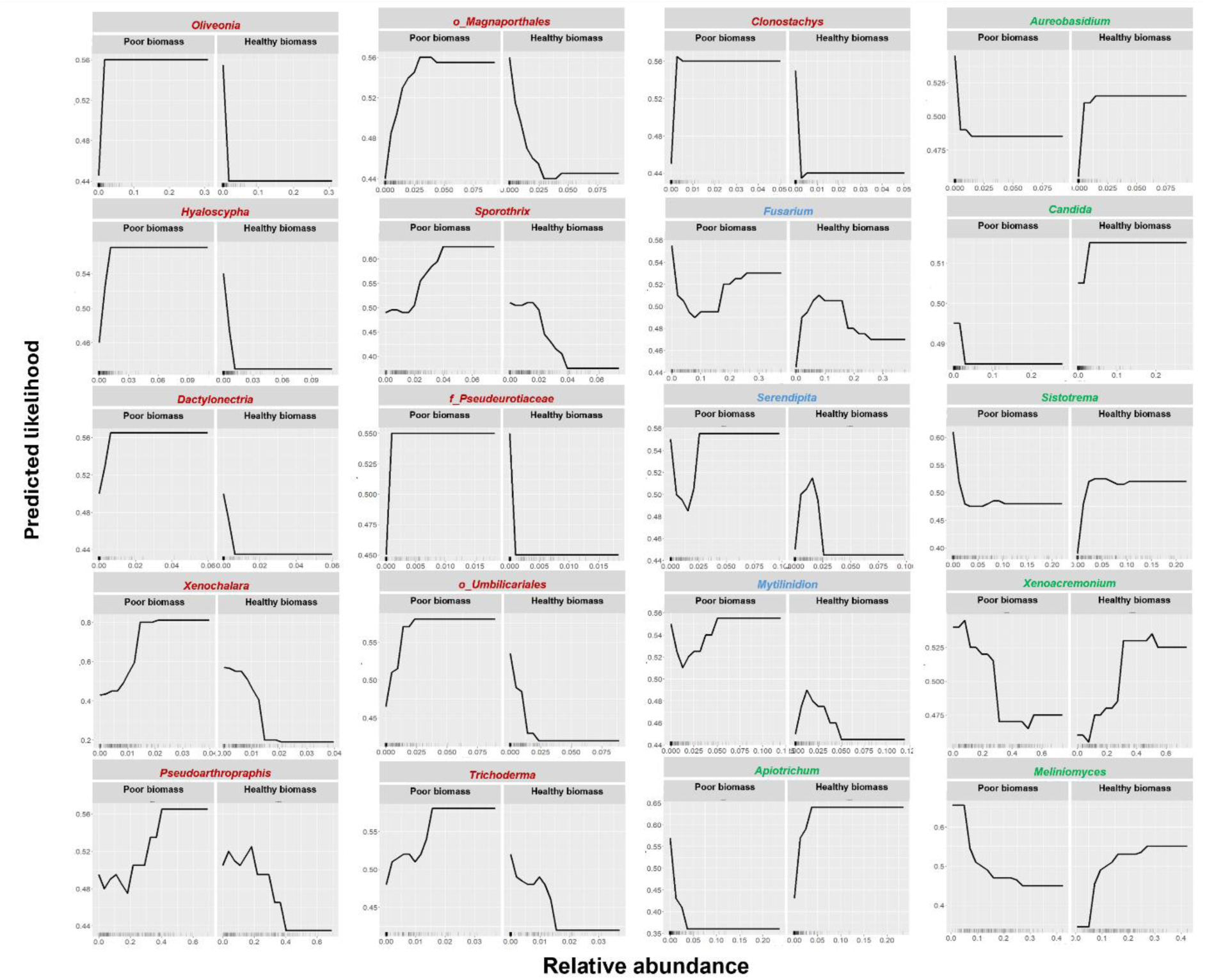
Top fungal features from GBM model predicting contributions to healthy vs. poor shoot biomass. Partial dependence plots (PDP) for the plot shows how changes in the abundance of a specific microbial genus influence the predicted likelihood of the growth condition across all samples. The plot visualizes how changes in the feature values influence the model’s prediction, holding all other variables constant. Color of the genera name indicates the contribution directions-Red = towards poor shoot biomass, Green= towards healthy shoot biomass, and blue = mixed contribution. Taxa names having prefix “o_” and “f_” indicate order or family as the respective genera could not be assigened.

## Discussion

Between the increasing demand for renewable wood products, and natural disasters such as forest fires, droughts, and flooding, there is growing demand for healthy, well adapted, cost-effective plantation seedlings for reforestation efforts. At the same time, traditional sources of inorganic fertilisers are becoming costly and unsustainable for large-scale production. In response, nurseries have begun to revise their growing protocols including varying fertiliser regimes. While past work has focused on altering inorganic nutrient inputs [78, 79], nursery managers are now attempting to meet growing demand for plantation seedlings by developing more sustainable procedures that learn from natural systems [80, 81]. Plantations in boreal and temperate zones are highly productive and their soils are typically characterised by a high level of organic nutrients that are released to plant roots via the activity of soil microbes. Therefore, there is large scope to incorporate more organic forms of nutrition to jump-start this microbial cycling in nursery settings with the aim to develop seedlings that are more robust when they are planted into plantation forestry sites. As opposed to our initial hypothesis, amino acid treatment appeared to drive pine root microbial assembly away from dependence on ectomycorrhizal fungi such as *Serendipita* and *Sistotrema* to a more diverse range of microbes that worked together in concert to promote plant health. Therefore, even in a well-studied systems such as radiata pine, there is still much to learn about their production in zones outside their native range.

Biostimulants have been on the market for many decades and include a range of different products from biological to chemical; all seek to improve plant growth either directly or indirectly. According to European Union regulations, biostimulants are not to be evaluated based on their nutrient content but, rather their ability to increase plant nutrient absorption and/or use efficiency, to improve abiotic stress, and to ameliorate produce quantity [82]. Some have gained widescale market approval and are therefore used consistently. Example include the use of protein hydrolysates [83, 84], seaweed extracts[85, 86], and silicon [87]. Our study supports the classification of amino acid fertiliser as a biostimulant as, compared to equivalent levels of inorganic nutrients, the amino acid applied significantly enhance the growth performance of radiata pine seedlings. Amino acid fertigation led to an increase in shoot biomass by >40%, an enhancement that aligns with previous studies where amino acid application promoted growth in various crops by improving nitrogen uptake and assimilation [29, 88]. While amino acids could serve directly as an available nitrogen sources, our study suggests it is more likely that they serve as signalling molecules that modulate physiological processes, including protein synthesis and hormonal regulation as previously reported [89, 90]. Interestingly, we observed a significant reduction in root biomass and root-to-shoot ratio in amino acid treated plants. This shift suggests that seedlings prioritize shoot growth over root development when amino acids are readily available, possibly due to enhanced nitrogen uptake efficiency reducing the need for extensive root systems. Similar observations and conclusions have been reported in other studies where improved nutrient availability led to reduced root growth but increased shoot biomass [31].

A critical aspect of our study is the alteration of the root fungal community in response to amino acid fertigation. We detected a significant increase in fungal diversity and richness, with distinct shifts in community composition compared to inorganic fertigation treatments. It was of interest to note that the known ectomycorrhizal fungi, which are typically associated with nutrient acquisition in pines [61, 62], exhibited a reduction in abundance. This would suggest that amino acid fertigation might reduce the plant’s reliance on ectomycorrhizal partners, shifting the microbial community toward beneficial non-mycorrhizal fungi. Of those fungi enhanced by amino acid treatment, *Meliniomyces* is known for its versatile lifestyle, functioning as an ericoid mycorrhizal, endophyte, and ectomycorrhizal fungus, all of which contribute to supporting plant growth [63]. *Xenoacremonium*, the most dominant genus in amino-acid-treated plants, remains poorly studied in terms of its role in plant biology. However, some studies suggest that this fungus may live as an endophyte and produce bioactive compounds [91]. Of those fungi repressed by amino acid treatment, *Dactylonectria* is the only genus within the group that is well-documented for causing root-associated diseases in plants, particularly conifers [74, 75]. *Xenochalara*, primarily known for its saprotrophic lifestyle, has also been reported to form consortia with pathogenic fungi, which can exacerbate plant diseases, as seen in black spruce [92]. *Sporothrix*, although better known as an animal pathogen, was found to be prevalent in the inorganic treatment and may negatively impact plant growth when present in high abundance [93]. The co-occurrence network analysis further illustrates the impact of fertigation type on not just individual genera/species, but also on fungal community structure. The amino acid fertigation network exhibits higher modularity and connectivity, indicating a more complex and interactive microbial community. Biostimulants have been found to have a range of effects on microbiome assembly and connectivity. Shi and colleagues [94] found that robustness of the mycobiome associating with maize was increased by only one of the biostimulants tested (a seaweed based product), and that this effect varied with plant genotype. Key hub genera were identified as *Apiotrichum, Oliveonia, Trichoderma, Hyalodendriella, and Trichomonascus* which indicate that they play central roles in maintaining network connectivity as has been found in other studies[95]. Interestingly, some dominant genera like *Xenoacremonium, Peudoarthrographis, Sistotrema, Meliniomyces,* and *Fusarium* either did not appear in the network or have low connectivity with other features. Other work in this field would suggest that, as highly connected microbial networks are considered robust [96], lack of connectivity of these microbes could put them at risk of loss should adverse environmental conditions be experienced. Therefore, while amino acid fertigation may have the dual effect of promoting beneficial fungi while suppressing pathogens to enhance plant resilience, loss/reduction of key ectomycorrhizal fungi due to this treatment should be considered and mechanisms derived to improve retention of these key fungi.

Amino acid fertigation also significantly altered the phytohormone profiles in radiata pine seedlings, potentially via the action of the microbiome. Within our dataset, *Apiotrichum*, *Aureobasidium*, *Candida* are known for their ability to promote plant growth mainly thorugh producing IAA [67, 97–99], an essential plant hormone that regulates cell elongation and division [99–101]. The increased levels of IAA may, therefore, have played a role in the higher shoot biomass seen in seedlings treated with amino acids. In contrast, amino acid treatment caused a significant reduction (approximately 50%) in the levels of gibberellins tested, specifically GA_3_ and GA_4_. Although GAs are, like IAA, associated with cell elongation and leaf expansion [102], IAA and ABA work antagonistically to GAs, creating a balance in growth regulation [103–105]. Thus, the induction of IAA accumulation in above-ground pine tissues may explain the repression of GA. We observe a weak, but statistically significant, negative relationship between GA_3_ and N. While our finding of a negative correlation of GA_3_ and total N content is contradictory with a previous study showing GA_3_ support nitrogen metabolism [106], the repression of GA_3_ may allow for higher N content and, by extension, enhanced chlorophyll content and photosynthetic efficiency. Amino acid fertigation increased total N levels in needles by approximately 50%, an increase that has previously been found to lead to higher chlorophyll content and photosynthetic productivity [28, 29]. Our observed improvements in Photosystem II (PSII) efficiency, as indicated by increased Fv/Fm and ΦPSII ratios, reflect a higher capacity of amino acid treated plants for light capture and energy conversion [58]. Additionally, positive effects on Photosystem I (PSI) parameters suggest enhanced electron transport capacity, supporting sustained photosynthetic activity. In contrast, plants treated with inorganic fertigation appeared to rely more on protective energy dissipation mechanisms, such as non-photochemical quenching (NPQ), indicating less efficient photochemical processes and a higher risk of photoinhibition [107, 108]. The higher NPQ values suggest that inorganic fertigation induces greater stress on the photosynthetic apparatus, possibly due to suboptimal nitrogen assimilation affecting chlorophyll synthesis and function. Taken together, our results indicate that amino acid addition to pine seedling roots has a system effect on needle chemistry and above-ground growth mediated through alterations within the mycobiome.

Given the complexity of microbiome data, especially when combined with plant growth factors, a number of machine learning techniques have been developed to identify key organisms within a microbiome that could be driving a plant phenotype of interest. Machine learning offers the advantage of accounting for non-linear relationships and complex interactions between microbial communities and plant traits, which traditional statistical methods might miss. Monte Carlo simulations have been used to understand microbial dynamics and shifts in medical studies [109], while Convolutional Neural Networks can be employed to capture unique microbial features within a dataset with high training accuracy[110, 111]. While the relative abundance and differential abundance analyses we undertook provided insights into specific genera enriched under different fertigation treatments, these methods alone do not capture the broader ecological interactions that contribute to plant health. We chose to use a gradient boosting model approach to our data analysis as this model sequentially corrects for errors rather than building multiple independent models which can lead to less accurate error correction [112]. Through the gradient boosting model, we were able to identify key microbial taxa whose presence or abundance is most predictive of healthy growth. Genera like *Apiotrichum, Aureobasidium, Candida, Sistotrema, Xenoacremonium,* and *Meliniomyces* positively contribute to healthy biomass accumulation. Conversely, taxa such as *Dactylonectria, Xenochalara,* and *Sporothrix* are associated with reduced shoot biomass, likely due to their pathogenic or saprotrophic lifestyles that can negatively impact plant health [75, 92]. The nuanced roles of genera like *Fusarium* and *Serendipita*, which show positive contributions at low abundance but negative effects at higher levels, highlight the complexity of plant-microbe interactions and the importance of microbial balance for optimal plant growth and new computational tools to understand these complex interactions. This approach, then, allows us to pinpoint microbial signatures of plant health and understand the cumulative effects of microbial diversity on growth outcomes, thereby providing a more holistic understanding of how microbial communities influence plant resilience and biomass accumulation. The identity of these keystone fungi could then be targetted in future for isolation and use in fungal inoculants, in concert with amino acid fertigation, to improve seedling health.

The ability of an amino acid biostimulant to enhance plant growth via the complex interplay between root-associated microbial communities, plant hormone profiles and photosynthetic efficiency presents an opportunity to develop integrated fertilization strategies that harness plant-microbe interactions for improved growth and resilience. Future research should focus on elucidating the specific mechanisms by which amino acid biostimulants influence microbial recruitment and activity with meta-transcriptomic approaches to provide deeper insights into microbial functions and their contributions to plant physiology. Furthermore, given the repressive role on pathogenic members of the mycobiome, work should look at the comparable resistance of biostimulant treated seedlings to external pathogens. Altogether, while our tests focused on controlled conditions in replica nursery containerised production, they demonstrate the promise that incorporation of biostimulants like amino acid fertigation into plantation forestry has on reducing inorganic fertilizer use and enhancing the sustainability of plantation forestry operations.

## Conclusion

Our work comparing equivalent total addition of inorganic versus organic N highlights the multifaceted benefits of amino acid fertigation in radiata pine seedlings, encompassing enhanced growth, optimized hormonal balance, and improved nitrogen uptake (Figure 9). Our findings also suggest that amino acid fertigation modulates the root microbiome to favour beneficial fungi that promote plant growth, although these are from genera that are typically not considered during design of growth-enhancing inoculum (Figure 9). These findings support growing evidence that amino-acid biostimulants represent an environmentally sustainable, effective alternative to inorganic nitrogen fertiliser, with significant forest nursery production advantages for species such as radiata pine. Future research should focus on long-term consequences of amino-acid fertigation on forest yield and ecosystem health, and on the optimization of formulations for application in forestry work on a broader scale.

**Figure 9:**
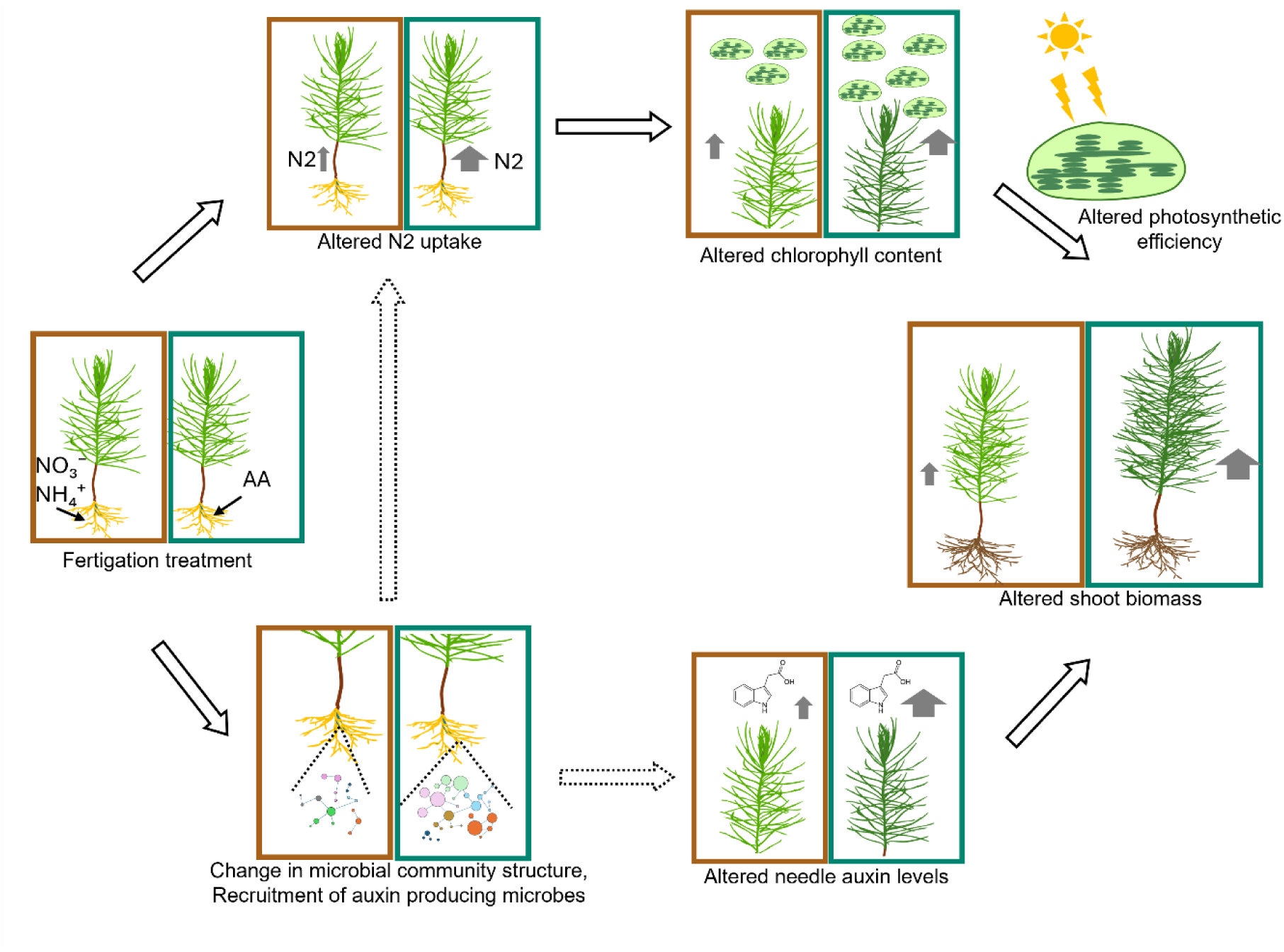
Schematic model showing how radiata pine seedlings, each receiving the same total nitrogen dose, diverge in growth responses according to whether the nitrogen source is inorganic or amino-acid-based. The layout begins (left) with identical nitrogen applications in either inorganic (red boxes) or amino-acid (green boxes) form. In seedlings treated with amino acids, enhanced nitrogen uptake increases chlorophyll content and photosynthetic capacity (top row), thereby boosting shoot biomass. Concurrently, amino-acid fertilization reshapes the root fungal community, favouring auxin-producing taxa that may elevate auxin levels in needles (bottom row) and further promote growth. The figure concludes (right) by underscoring these two synergistic pathways through which amino-acid fertilization surpasses inorganic treatments in promoting radiata pine performance.

## Declarations

### Ethics approval and consent to participate

Not applicable. This study did not involve human participants or animal experiments requiring ethical approval.

### Consent for publication

Not applicable. The manuscript does not contain any individual person’s data in any form.

### Funding

This research was supported by the National Institute for Forest Products Innovation (NIFPI) program funded by the Australian Government Department of Agriculture, Fisheries and Forestry and the Victorian Government, and 25 large forest management entities through Forest and Wood Products Australia (FWPA) under the project titled ‘Innovative nursery management solutions to sustainably manage root disease, improve nursery utilization, and enhance resilience and productivity of planted pines’ (NV070).

### Availability of data and materials

The raw Illumina MiSeq ITS2 reads generated in this study are available in the NCBI Sequence Read Archive under BioProject PRJNA1267575. (https://www.ncbi.nlm.nih.gov/sra/PRJNA1267575)

### Competing interests

The authors declare that they have no competing interests.

### Author contributions

All authors contributed to the conception and study design, JC and BT performed the experiment and data collection. JC, RS, KLP and JMP analysed and interpreted the data. JC and JMP drafted the manuscript and all authors contributed to editing and revising the manuscript.

## Acknowledgement

We acknowledge the significant financial contributions from industry members and NIFPI’s commitment to advancing Australia’s forest and wood products industry through innovation in areas such as forest management, timber processing, and the bio-economy.

## Supporting information

**Figure S1: Principal Component Analysis (PCA) biplot of morphological and photosynthetic variables measured under four different fertiliser treatments.**

Dots represent individual samples, colored and enclosed by ellipses indicating the 95% confidence region for each fertiliser group. The PC1 and the PC2together explain 34.58 % and 25.02 % of the total variance, respectively. The vector’s direction and length indicate the traits’ contribution to the first two components in the PCA. Morphological variables include shoot weight (SW_avg), plant height (PH_avg), collar diameter (CD_avg), and root–shoot ratio (Rw_SW_ratio), while photosynthetic traits include total chlorophyll (total_chl), chlorophyll a (Chl_a), chlorophyll b (Chl_b), and PSI activity measurements.

**Table S1:** Genus/species used as microbial inoculant

**Table S2:** Effect of fertigation on Radiata pine Growth Morphology

**Table S3:** Effect of fertigation on Radiata pine needle chlorophyll and total N content

**Table S4:** Effect of fertigation on Radiata pine Photosynthetic Efficacy

**Table S5:** Effect of fertigation on Radiata pine needle hormone level

**Table S6:** Pearson’s correlations of needle hormone levels with major root fungal genera found in the pine root

## Notes

### Competing Interest Statement

The authors have declared no competing interest.

### Summary of Updates

Minor changes in the abstract and figure number.

