## Supplementary material for "Amino acid biostimulant increases radiata pine photosynthetic efficiency and growth through optimised mycobiome and nitrogen assimilation": Fig S1 and S2

**Figure S1**

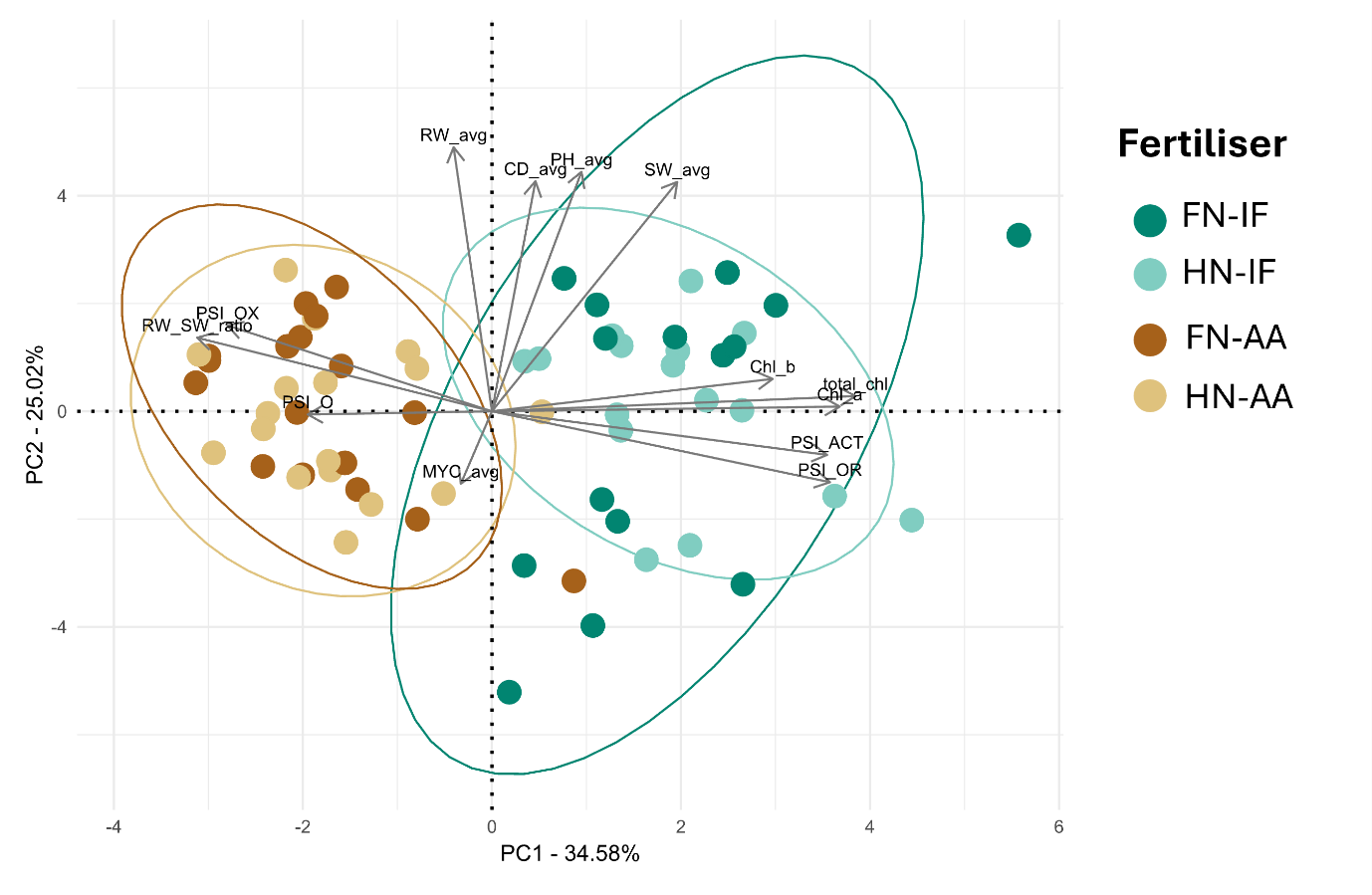

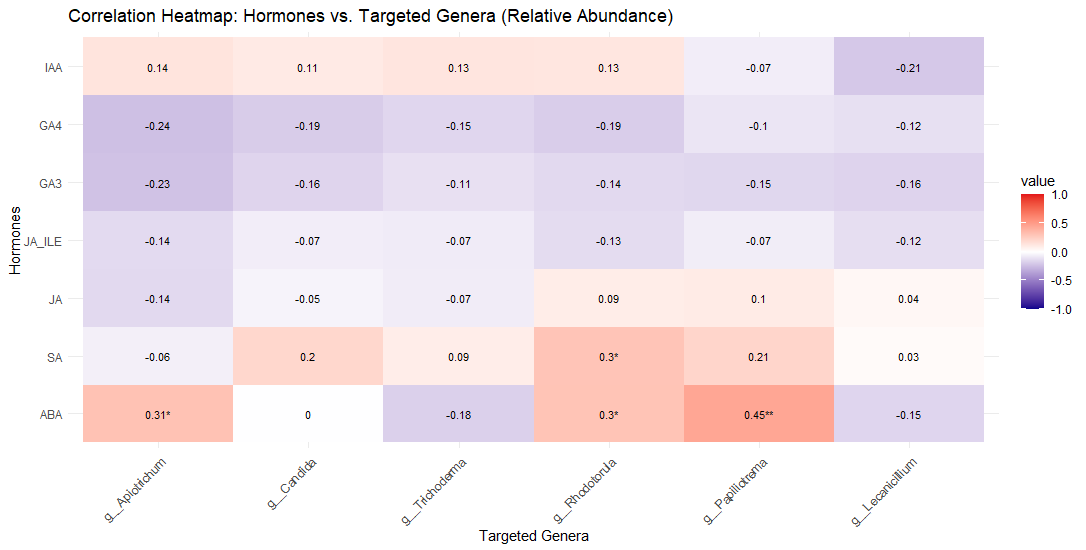

**FigS2** Correlation matrix between the targeted auxin producing genera in the root and phytohormone level in needles. The color gradient ranges from blue (strong negative correlation) to red (strong positive correlation). Numeric values in each cell represent Pearson’s r, whereas * denotes p < 0.05 and ** denotes p < 0.01.
